# Stable fusion of bone marrow-derived cells with Purkinje neurons enables long-term nuclear integration and neuronal repair

**DOI:** 10.64898/2025.12.31.694485

**Authors:** Elisabeth M. M. Meyer, Siân Baker, Benjamin Thewlis, Nicole Jastrzebowska, Bruno Salomone Gonzalez De Castejon, Paul Chadderton, Kevin C. Kemp

**Affiliations:** School of Psychology and Neuroscience, University of Bristol, UK

## Abstract

Developing effective strategies to replace or restore injured neurons could significantly revolutionise the treatment of currently incurable neurodegenerative disorders. Studies have shown that bone marrow-derived cells (BMDCs) can migrate into the brain and fuse with damaged Purkinje neurons (PNs), restoring their structure and function. Yet, despite over two decades of research, the temporal dynamics and functional stability of these fusion events *in vivo* remain poorly defined, largely due to reliance on static, post-mortem histological analyses that provide only a single snapshot in time. Here, using *in vivo* two-photon imaging of the intact cerebellum in EGFP-bone marrow chimeric mice, we visualised the dynamics of BMDC-PN interactions both in real time and longitudinally to monitor the physiological characteristics of fused PNs. Notably, we provide definitive evidence that fused cells are not transient but remain stable over extended periods in the living brain. Moreover, we demonstrate sustained transcriptional activity of donor-derived genes within fused PNs, indicating partial or complete functional integration of the BMDC nucleus. Given their accessibility and genetic tractability, BMDCs represent a practical and promising platform for targeted delivery of therapeutic genes to PNs, opening new avenues for treating neurodegenerative disease.

## Introduction

The cerebellum or ‘little brain’ plays a critical role in control and coordination of movements in addition to higher-order functions including cognition, memory, and emotional processing. Cerebellar injury occurs relatively commonly in children and adults around the world^1^ via numerous mechanisms including genetic disease, stroke, metabolic disturbances, infection, autoimmune disease, and cancer^2^ causing loss of Purkinje neurons (PNs). PNs provide the only output of the cerebellar cortex and are not generated after birth^3^, as such, their loss is characterised by chronic ataxia, impaired muscle coordination, weakness, difficulties with speech, swallowing and eye movements. Developing effective treatments aimed at replacing or restoring injured PNs could prove highly impactful across a wide range of disorders.

Experimental findings in both humans and rodent models of neurodegenerative disease have shown that cells derived from the bone marrow (BM) can migrate into the brain and merge (fuse) with damaged PNs^3-7^. During fusion, the BM-derived cell (BMDC) transfers a healthy nucleus to the damaged neuron leading to repair and restoring the ability to process and transmit information^8^. This natural phenomenon offers a promising pathway to overcome the difficulty of replacing injured neurons.

Cell and gene therapies offer real hope for cerebellar repair, but fundamental gaps remain in our understandings of cell fusion in the CNS. Despite over two decades of study, we know very little about how cell fusion is controlled or its physiological role in the cerebellum and elsewhere. In particular, little is known about constitutive rates of fusion *in vivo*, and the stability of donor-recipient neurons over time. By applying recent advances in genetic engineering and longitudinal *in vivo* imaging, we aimed to investigate, for the first time, the frequency of fusion and stability of fused PNs in the live cerebellum. Tracking cell fusion and fused PNs over prolonged periods in the intact cerebellum will be critical for advancing this process towards a regenerative therapy for clinical application.

## Materials and methods

### Animals and experimental design

All animal experiments were performed in accordance with the UK Animals (Scientific Procedures) Act 1986 and approved by the University of Bristol Animal Welfare and Ethical Review Body. The study protocol was developed in accordance with the ARRIVE guidelines using methodologies validated in our previous published studies and protocol papers^8-10^. Transgenic mice ubiquitously expressing EGFP (strain C57BL/6-Tg(CAG-EGFP)131Osb/LeySopJ, stock # 006567) and those expressing Cre recombinase under the control of the L7-Pcp2 promoter (strain B6.129-Tg(Pcp2-cre)2Mpin/J, stock # 004146) [L7Cre-2 mice] were obtained from Jackson Laboratory, US. All mice were group-housed at the University of Bristol’s Animal Services Unit (ASU) with environmental enrichment in a pathogen-free facility, with free access to food and water. Post bone marrow transplant (BMT), mice were maintained in filter top cages (pore size 100 μm) and weighed daily (for a minimum 2 weeks) during the recovery period. The study followed a single-group exploratory design. Given the novelty of the study, a small sample size was used to gather data on PN fusion frequency/stability to inform future work. No unexpected adverse effects were reported.

### Bone marrow transplantation: generation of EGFP-expressing BM chimeric mice

Donor BM cells were harvested, under sterile conditions, from 10 - 12-week-old male/female C57BL/6 EGFP-expressing transgenic mice^8,9^. Recipient L7Cre-2 male and female mice (aged 12 weeks) were irradiated, with a single dose of 950 rad from a 137Cs source, 6 hours prior to receiving ≥1 x 10^7^ unfractionated EGFP-expressing BM cells in PBS by tail-vein injection. Sterile water, antibiotics (Baytril), sterile food and bedding were provided for 6-weeks post-transplant.

Detection of EGFP chimerism in peripheral blood was performed at 6 weeks post BMT (animals aged 18 weeks) by flow cytometry (NovoCyte flow cytometer, Agilent) as previously described^8^. Data were evaluated using NovoExpress software.

### Cranial window surgeries

Virus injections into the vermis and cranial window implantation were performed as previously described in BMT L7Cre-2 mice^11^ to transduce cerebellar PNs with the cre-dependent fluorophore tdTomato. Briefly, BMT mice were anaesthetized using Isoflurane (3-5% induction, 1.5% maintenance). Carprofen (5 mg/kg) and Buprenorphene (0.075 mg/kg) were injected subcutaneously. Lidocaine (2mg/kg) was injected locally on the scalp and muscles attached to the occipital ridge (m. semispinalis capitis). The skin above the skull was removed and the muscles gently pushed aside to expose the location of the craniotomy (central vermis, ∼3 mm). 5 viral injections (virus: AAV(1)-CAG-Flex-tdTomato, Dr. Ed Boyden, UNC vector core) were performed at ∼300um depth at various locations within the craniotomy (∼150nl per injection), and a 3mm diameter glass coverslip was fitted over the craniotomy. Skull and muscle tissue was covered in Histoacryl (Braun) and a custom designed headplate was fixed to the top of the skull^12^ using dental cement (Paladur). Animals were continuously group-housed (3-4 per cage) during surgery recovery (∼21 days) and imaging.

### Two-photon imaging

For anaesthetized two-photon imaging (used for all video imaging), animals were injected intraperitoneally with 0.05 mg/kg Fentanyl, 5 mg/kg Midazolam and 0.5 mg/kg Medetomidin in sterile saline (0.9% NaCl). Once animals had fallen asleep, they were head-fixed to a custom-built stage, equipped with a body temperature monitor. For awake two-photon imaging (used for all longitudinal imaging), animals were head-fixed on a custom-built stage under the 2-photon microscope (Hyperscope, Scientifica) equipped with a galvanometer for x- and y-axis imaging respectively (Cambridge Technologies, 8315KL). The angle of the objective (16x, NA 0.80, Nikon) was set to 30 degrees for all experiments (vertical: 0 degrees). Fluorophores were visualised using a femtosecond laser (Chameleon Ti:Sapphire, Coherent) tuned to 930 nm. Z-stacks of neurons and BMDCs in the cerebellar vermis were acquired at a rate of 0.3 to 0.7 frames/s for 512 x 512 pixels, with a distance of 2um between planes, and a minimum of 4 frames/plane. Each plane was later averaged to improve the signal to noise ratio and compensate for potential movement artefacts. Time series were acquired at a rate of 0.4 frames/sec. For chronic imaging experiments, laser power and detector sensitivity (PMT) levels were kept constant between experiments to allow for fluorescence comparisons across experiments. To identify previously recorded PNs, a map of the blood vessels and imaging site was created using the ‘Position Save Module’ plugin in SciScan (Scientifica). This plugin recorded locations (extracted coordinates x, y and z of the micromanipulators moving the objective) along with corresponding screen shots of their fields of view. PNs were then re-identified based on the relative distance of the imaging site to landmarked blood vessels.

### Fluorescence intensity measurements

Two-photon image stacks were processed using ImageJ (NIH). BMDCs were identified based on their green fluorescence. Fused cells were identified based on overlapping red and green fluorescence, characteristic PN morphology, and localisation within the Purkinje cell layer. For each fused PN, the soma was manually outlined in three z-planes corresponding to maximal cross-sectional area. Mean fluorescence intensity was measured within each region of interest (ROI) and normalised by subtracting the mean EGFP intensity to three EGFP-positive BMDCs within the same imaging field. This normalisation controlled for inter-session variability in window clarity.

### Data analysis

Statistical tests were performed using GraphPad Prism (v. 10.5, GraphPad Software Inc, USA). Values of P < 0.05 were considered statistically significant. Where data were known or predicted to violate assumptions for parametric statistical testing (normal distribution or homogeneity of variance), an equivalent non-parametric test was performed. Fused PNs were identified and counted by eye.

### Data availability

All data that support the findings of this study are available within the article and figures. Raw microscopy data are available from the corresponding author, upon reasonable request.

## Results

### Bone marrow chimera formation and window implantation

To track BM-derived cells infiltrating the cerebellum, fifteen EGFP-BM chimeras were generated by exposing L7Cre-2 mice to myeloablative irradiation, followed by stable reconstitution with EGFP-expressing BM cells harvested from transgenic donor mice (**Figure 1A**). All chimeric mice showed robust levels of chimerism; 89 ± 1.9 % (*n* = 15) of mononuclear cells within the peripheral blood of transplanted mice were EGFP-positive and thus donor-derived. Once chimerism was established, cranial windows were implanted over the cerebellar vermis following injections of AAV(1)-CAG-Flex-tdTomato to label PNs at the window site (**Figure 1B**).

**Figure 1.**
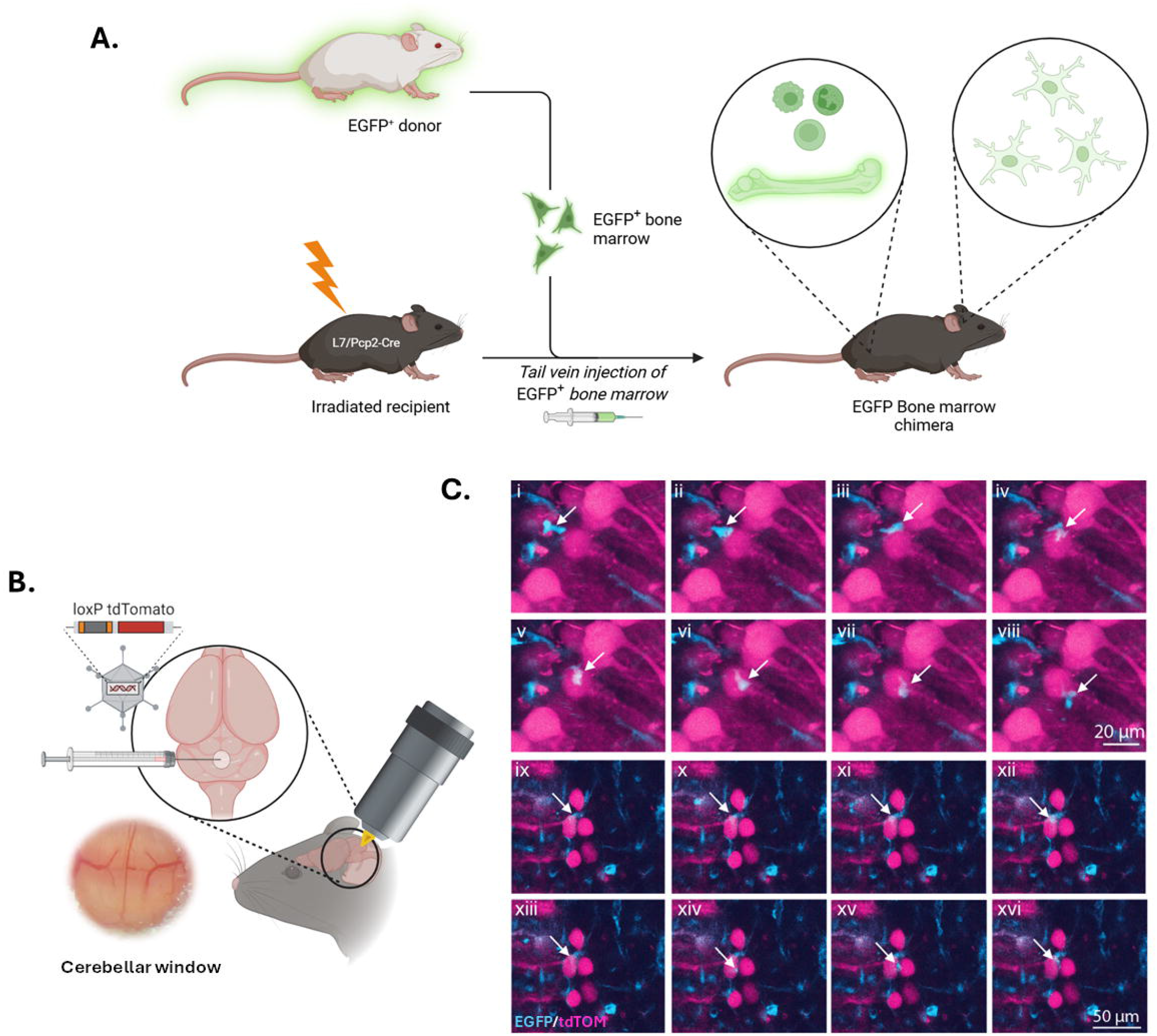
EGFP-BM chimerism coupled with two-photon microscopy can be used to successfully monitor BMDC temporal dynamics in the live intact cerebellum. Schematic representations of the experimental design. **A**. To form EGFP-expressing BM chimeras, recipient L7Cre-2 mice underwent myeloablative irradiation prior to receiving unfractionated BM cells expressing EGFP by tail-vein injection. **B**. Following successful immune engraftment, mice underwent craniotomy, localised injection of a Cre-dependent AAV encoding tdTomato, cranial window implantation, and subsequent two-photon imaging. **C**. Two-photon imaging of the cerebellum showing highly motile EGFP-BM-derived cells (cyan) interacting with PNs (magenta) in real-time (Representative examples from 2 animals. I to VIII: elapsed time between frames: 19 s. IX to XVI: time between frames: 15.2 s).

### *In vivo* two photon imaging of fused Purkinje neurons

Regions with high levels of red fluorescence (reflecting tdTomato expression) were chosen as imaging sites. Using both z-stacks and time series acquisition (see Methods), we were able to track the dynamics of EGFP-BM cell-neuron interactions in real time and assess number of fused cells. BMDCs were shown to adopt a ramified microglial-like morphology, and, being highly motile within the brain, shown to visit, and contact, multiple PNs within individual imaging sessions (**Figure 1C, Supplementary Video 1, Supplementary Video 2**). Sustained envelopment of the PN soma by EGFP-expressing cells was observed in several mice **(Figure 2A, Supplementary Video 3)**.

**Figure 2.**
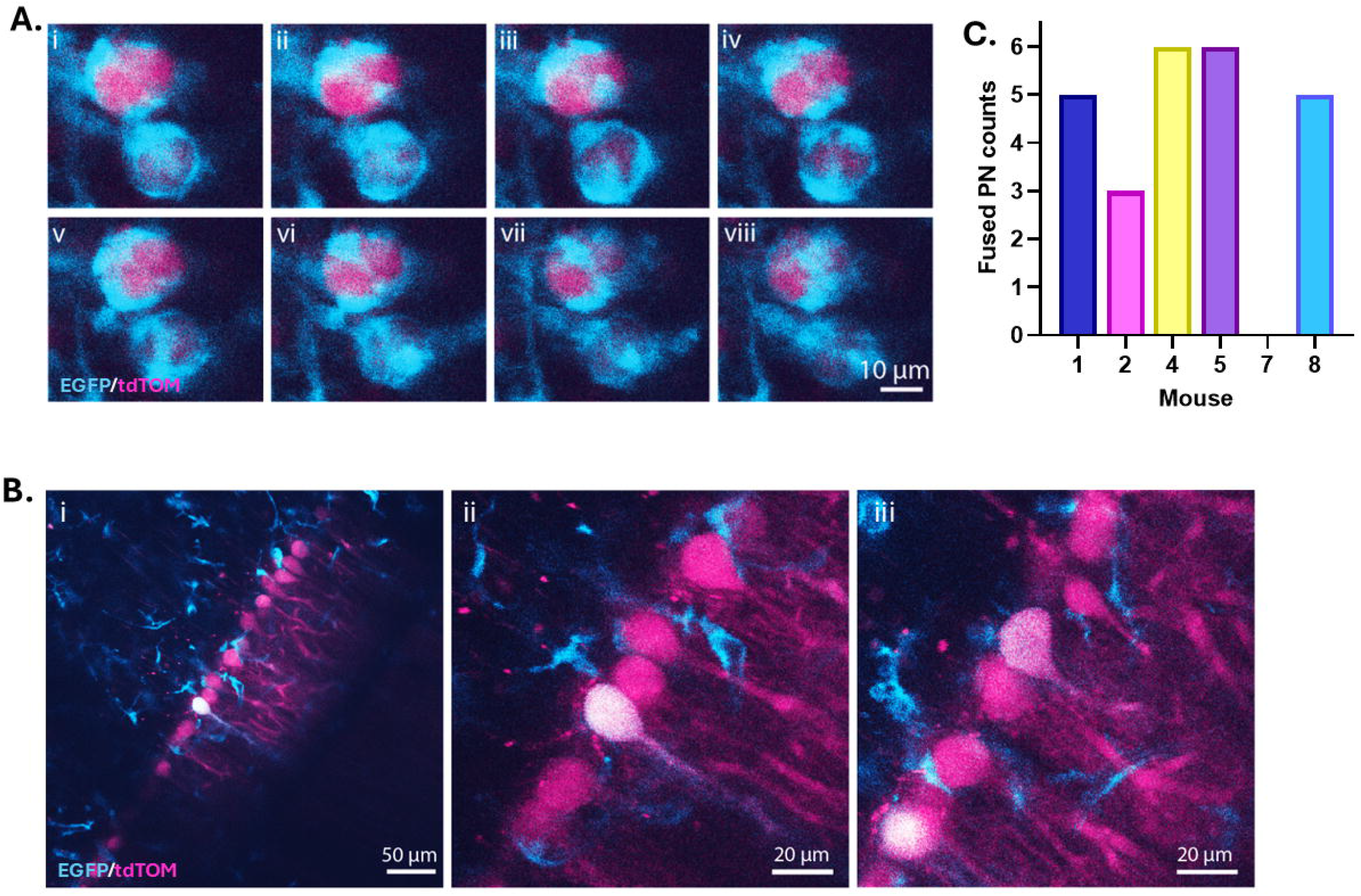
Fused EGFP-expressing PN are detected *in vivo* using two-photon imaging. **(A)** Envelopment of the PN soma (magenta) by EGFP-expressing cells (cyan) in real time (time between frames: 22.8 s). **(B)** Representative example of fused PNs at a single imaging location and **(C)** quantification of fused EGFP-expressing PN that could be observed within the entire cerebellar window of each animal, *n* = 6 animals (5 animals: 4 weeks of imaging, 1 animal: 8 weeks of imaging), n = 3 animals [mouse IDs 3, 6, and 9]: no fused cells could be observed at last imaging timepoint due to the window being overgrown by scar tissue (2 animals: 4 weeks of imaging, 1 animal: 8 weeks of imaging). Only animals which could be imaged at the last imaging time point are included in this figure.).

Dual expression of tdTomato and EGFP was used to identify PNs that had fused with EGFP-expressing BMDCs; EGFP expression confirming donation of BM-derived nuclear transgenes within the host PN. *In vivo* imaging revealed fusion events were evenly scattered throughout the vermis in most mice examined within the study (11/15 animals) (**Table 1, Figure 2B, 2C, Supplementary Video 4**). EGFP-positive PNs exhibited the typical, well-defined PN morphology, characterised by extensive dendritic arborisation climbing through the molecular layer (**Figure 2B**).

**Table 1.** Animals used for in-vivo two-photon imaging.

| **Total number (n = 15)** | **Single**<br>**imaging** | **4 weeks**<br>**imaging** | **8 weeks**<br>**imaging** |
| --- | --- | --- | --- |
| **Number of animals for single imaging session** | 6 | - | - |
| - *No fusion detected during imaging* | 1 |  |  |
| **Number of animals for longitudinal imaging** | - | 7 | 2 |
| **Imageable for whole experiment** |  | 5 | 1 |
| - *No fusion detected during imaging duration* |  | 1 | 0 |
| **Window overgrown before end of experiment** |  | 2 | 1 |
| - *No fusion visible before window became overgrown* |  | 1 | 1 |

### Long-term stability

It is unknown if fused PNs are a transient phenomenon^19^ or binucleate fused cells represent a transient phase prior to complete ‘takeover’ by the fusing cells, leading to mononucleation^5,19^. By returning to the same regions across multiple sessions, we longitudinally monitored the stability and dynamics of fused PNs in 9 of the 15 animals for 4 or 8 weeks by recording the number and position of fused PNs at weekly intervals (see Methods).

EGFP-positive PNs were observed in the live brain as early as 2 months post-transplant. In animals with imageable windows throughout the whole experiment, the total number of EGFP-positive PNs that could be observed within the window site after 3 months (7-weeks post implantation of cranial window) was 4.2 ± 1.0 (*n* = 6 animals).

Remarkably, all fused cells remained detectable for successive weekly imaging sessions over a 4-week period (maximum 5 imaging sessions per mouse) (**Figure 3A, 3B)**. The maximum number of fused cells observed through the window after four weeks was six. In a single animal, we identified several fused PNs in week one and show that fused PNs were stable for at least two months (**Figure 3B)**. Cell fusion in the cerebellum thus occurs constitutively and is stable within the population.

**Figure 3.**
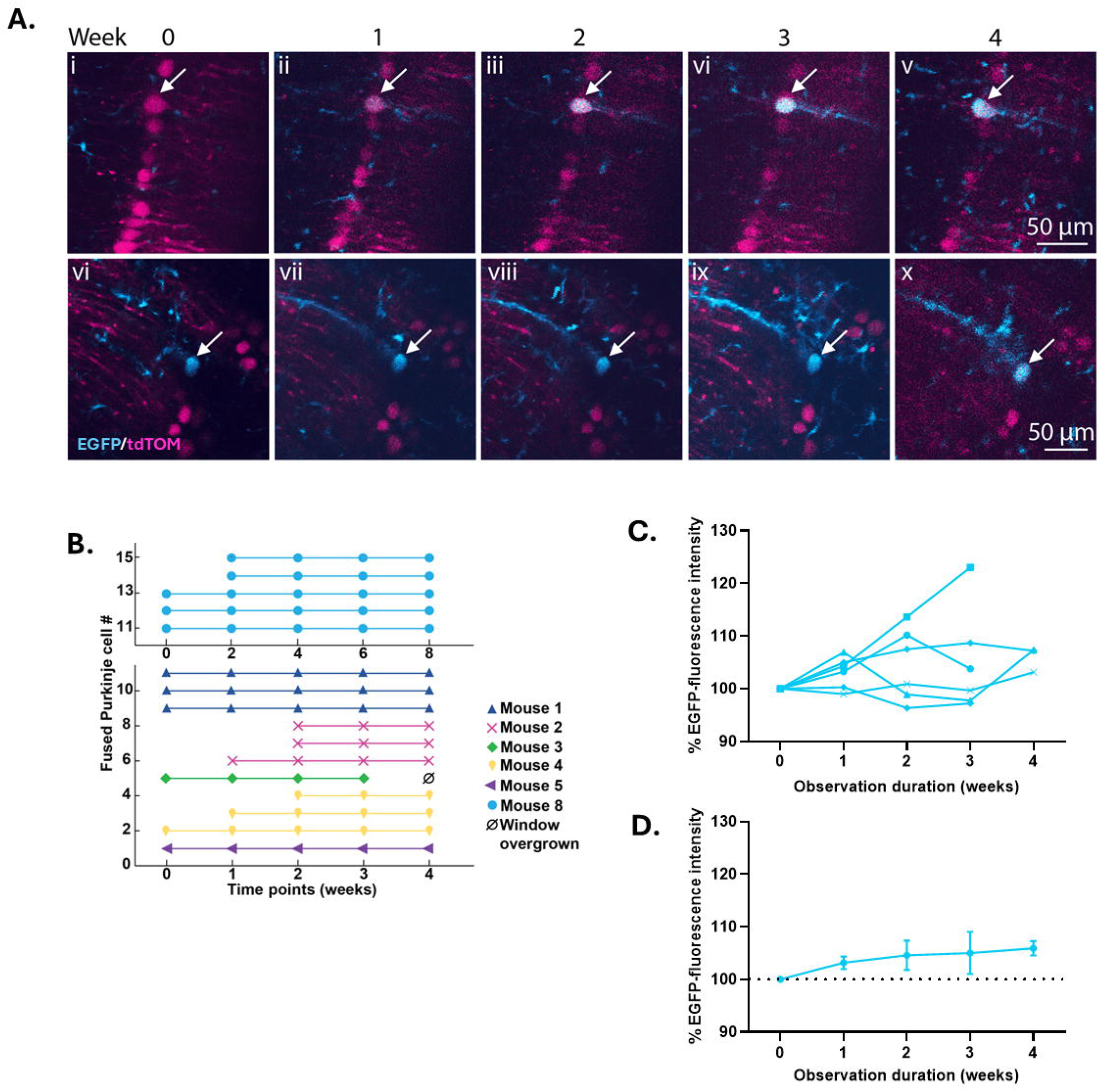
Longitudinal monitoring across multiple imaging sessions reveals fused PNs express donated genes for at least 2 months. **(A)** Longitudinal monitoring of two distinct cerebellar regions over a 4-week period, showing the appearance of a newly formed fused EGFP-expressing PN (cyan) at week 2 (i-v), and a fused PN (cyan) present for the entire study (vi-x). **(B)** A schematic showing the week fused PNs were identified and their presence in subsequent weeks. Mice were monitored over a 4- or 8-week period (4 weeks: n = 7, 8 weeks: n = 2. Only animals with identified fused PNs are included in this figure; no fused PNs found in mouse IDs 6 and 9). Weekly EGFP-fluorescence intensity levels of **(C)** individual and **(D)** all PNs post identification (mean ± SEM, *n* = 6 PN, Kruskal-Wallis test comparing weeks; P = 0.44).

### Transgene activation and expression in fused Purkinje neurons

Whether the donated and endogenous nuclei persist in expressing its own nuclear programme or one programme dominates post-fusion^5^ is largely unknown. For six fused PN, we used EGFP fluorescence intensity as a quantitative reporter of transgene mRNA expression in BM-derived nuclei transferred to the PN soma. In a newly identified fused PN (absent at timepoint 1), EGFP levels appeared to increase progressively over a 2-3 week period (**Figure 3B, C**). Across all fused PNs, sustained EGFP expression was observed over the 4-week study period, with the lower limit of standard error for changes in EGFP expression intensity remaining above the null value of 100% for the entirety of the study (**Figure 3D**). Statistically, there was no significant change in EGFP intensity over time (*P* = 0.44).

## Discussion

This study demonstrates definitively that BMDCs fuse with cerebellar neurons, resulting in stable, long-term integration of BM nuclear material into the PN population. In contrast to previous seminal studies that have relied solely on snapshot post-mortem histological analysis^3,5,6,8,13-17^, our longitudinal *in vivo* imaging provides a temporally resolved view of BM-PN fusion progression in the intact cerebellum. Using BM chimeric mice under physiological conditions, we show cell fusion occurs in the healthy cerebellum. A potential homeostatic role may be proposed, whereby BMDCs provide restorative cellular machinery and/or genetic material to PNs during adult life.

Importantly, our data demonstrate unequivocally, for the first time, that fused cells are not merely a transient phenomenon but remain stable for prolonged periods. All fused PNs persisted throughout the entire duration of the experiment. They continually expressed BM-derived donor genes, evidenced by sustained EGFP expression for at least two months, with no detectable decline in signal intensity. Expression of the donated genes incorporated into the host transcriptome is likely driven by exposure to transcription factors present in the PN. Indeed, the short half-life of EGFP excludes direct protein transfer as a source of expression^18^, and transfer of mRNA^19^ is unlikely in this model given poor stability and cytoplasmic degradation. Fascinatingly, in a single newly identified fused cell we were able to observe EGFP expression progressively increasing over several weeks, providing critical insights into the temporal dynamics of donor gene translation.

Using sex-mismatched transplantation, we have previously demonstrated in this model, and others in human tissue, that fused PNs form binucleate heterokaryons containing a complement of nuclei consistent with fusion between a PN and incoming BMDC^3,8^. Indirect evidence had previously suggested that these heterokaryons were stable^14,6^, however, others have argued that fusion does not yield stable heterokaryons under non□invasive conditions^15^. While heterokaryon instability was not explored in this study, stable transcriptional activity of the (donor) EGFP gene indicates partial, or even complete, functional integration of the incoming BMDC nucleus.

Fusion events continued to be observed up to 6 months following transplantation, and up to 11 weeks after cerebellar window implantation. These findings may suggest that cell fusion is an ongoing process rather than an immediate response or artifact of CNS injury. While we cannot exclude these events being caused by the long-term effects of BM-transplantation or cerebellar window implantation, previous studies have shown that fusion can occur in the complete absence of BM transplantation, radiation, or direct injury to the cerebellum^6,15,20,21^. Although only up to six fused cells were detected at the cranial window site, it is important to note that this region represents only a small fraction of the entire cerebellar cortex. Previous studies have reported low total numbers of fused cells across the whole cerebellum at 5-9 months post-BM transplantation^5,6,21^. However, *in vivo* two-photon imaging likely provides a more accurate assessment of fusion events, as it avoids several significant limitations including antibody labelling, sequential tissue sectioning, and section thickness (particularly relevant when monitoring fusion by PN binucleation) that all likely result in an underestimation of fusion events.

In the healthy brain, PN fusion occurs at low levels, yet striking increases (10 to 100-fold) in fusion are observed following injury and inflammation^5,6,8,22^. Previous studies showed up to ∼5% of all PNs fused in mice^6^, equivalent to ∼ 1 million cells in the human brain^23,24^. In patients with neurological disease, elevated numbers of binucleated PNs have been observed^7,20^, and we showed that fusion can repair aberrant electrical activity of injured PNs^8^. This natural mechanism of neuronal repair holds considerable promise for treating neurodegenerative and autoimmune conditions effected by PN loss or dysfunction^25^. Importantly, we have also shown evidence for fusion between BMDCs and large sensory neurons of the dorsal root ganglion^23^, highlighting broader potential for neuronal regeneration and gene delivery strategies. Given the accessibility and modifiability of BM cells, they represent a practical and promising vehicle for delivering therapeutic or corrective genes to PNs and other neurons across the nervous system.

## Acknowledgements

We gratefully acknowledge Dr. Paul Anastasiades and his research group for providing L7Cre-2 mice.

## Author contributions

EMMM, SB, PC and KCK planned experiments and interpreted data. SB, BT, NJ, BS and KCK performed bone marrow transplants. EMMM performed cranial window surgery and imaging experiments. EMMM, SB and BT performed data analysis. EMMM, SB, BT, PC and KCK wrote the manuscript.

---

**Supplementary Video 1. Time-lapse video of BMDCs moving and interacting with PNs in the cerebellum**. Video recorded at 0.4 frames/s and shown at 10 frames/s. Magenta: PNs. Cyan: BMDCs.

**Supplementary Video 2. Time-lapse video of BMDCs moving and interacting with PNs in the cerebellum**. Video recorded at 0.4 frames/s and shown at 10 frames/s. Magenta: PNs. Cyan: BMDCs.

**Supplementary Video 3. Time-lapse video of BMDCs enveloping individual PNs in the cerebellum**. Video recorded at 0.4 frames/s and shown at 10 frames/s. Magenta: PNs. Cyan: BMDCs.

**Supplementary Video 4. Z-stack of fused and non-fused PNs in the cerebellum**. Video showing scan beginning with superficial PNs extending 312 µm (78 images) deep into the cerebellum. Magenta: PNs. Cyan: BMDCs. White: double-positive PNs (fused PNs).

## References

1. Salman MS. Epidemiology of Cerebellar Diseases and Therapeutic Approaches. Cerebellum. Feb 2018;17(1):4–11. doi:10.1007/s12311-017-0885-2

2. Ashizawa T, Xia G. Ataxia. Continuum (Minneap Minn). Aug 2016;22(4 Movement Disorders):1208–26. doi:10.1212/CON.0000000000000362

3. Weimann JM, Charlton CA, Brazelton TR, Hackman RC, Blau HM. Contribution of transplanted bone marrow cells to Purkinje neurons in human adult brains. Proc Natl Acad Sci U S A. Feb 18 2003;100(4):2088–93. doi:10.1073/pnas.0337659100

4. Kemp K, Wilkins A, Scolding N. Cell fusion in the brain: two cells forward, one cell back. Acta Neuropathol. Nov 2014;128(5):629–38. doi:10.1007/s00401-014-1303-1

5. Johansson CB, Youssef S, Koleckar K, et al. Extensive fusion of haematopoietic cells with Purkinje neurons in response to chronic inflammation. Nat Cell Biol. May 2008;10(5):575–83. doi:10.1038/ncb1720

6. Magrassi L, Grimaldi P, Ibatici A, et al. Induction and survival of binucleated Purkinje neurons by selective damage and aging. J Neurosci. Sep 12 2007;27(37):9885–92. doi:10.1523/JNEUROSCI.2539-07.2007

7. Kemp KC, Cook AJ, Redondo J, Kurian KM, Scolding NJ, Wilkins A. Purkinje cell injury, structural plasticity and fusion in patients with Friedreich’s ataxia. Acta Neuropathol Commun. May 23 2016;4(1):53. doi:10.1186/s40478-016-0326-3

8. Kemp KC, Dey R, Verhagen J, Scolding NJ, Usowicz MM, Wilkins A. Aberrant cerebellar Purkinje cell function repaired in vivo by fusion with infiltrating bone marrow-derived cells. Acta Neuropathol. Jun 2018;135(6):907–921. doi:10.1007/s00401-018-1833-z

9. Kemp K, Hares K. Analyzing cell fusion events within the central nervous system using bone marrow chimerism. Methods Mol Biol. 2015;1313:165–84. doi:10.1007/978-1-4939-2703-6_12

10. Meyer EMM, Chadderton P. Tactile stimulation transiently disrupts encoding of whisker position by cerebellar molecular layer interneuron ensembles. bioRxiv. 2025:2025.08.12.669821. doi:10.1101/2025.08.12.669821

11. Leinweber M, Zmarz P, Buchmann P, et al. Two-photon calcium imaging in mice navigating a virtual reality environment. J Vis Exp. Feb 20 2014;(84):e50885. doi:10.3791/50885

12. Barkus C, Bergmann C, Branco T, et al. Refinements to rodent head fixation and fluid/food control for neuroscience. J Neurosci Methods. Nov 1 2022;381:109705. doi:10.1016/j.jneumeth.2022.109705

13. Alvarez-Dolado M, Pardal R, Garcia-Verdugo JM, et al. Fusion of bone-marrow-derived cells with Purkinje neurons, cardiomyocytes and hepatocytes. Nature. Oct 30 2003;425(6961):968–73. doi:10.1038/nature02069

14. Weimann JM, Johansson CB, Trejo A, Blau HM. Stable reprogrammed heterokaryons form spontaneously in Purkinje neurons after bone marrow transplant. Nat Cell Biol. Nov 2003;5(11):959–66. doi:10.1038/ncb1053

15. Nern C, Wolff I, Macas J, et al. Fusion of hematopoietic cells with Purkinje neurons does not lead to stable heterokaryon formation under noninvasive conditions. J Neurosci. Mar 25 2009;29(12):3799–807. doi:10.1523/JNEUROSCI.5848-08.2009

16. Nygren JM, Liuba K, Breitbach M, et al. Myeloid and lymphoid contribution to nonhaematopoietic lineages through irradiation-induced heterotypic cell fusion. Nat Cell Biol. May 2008;10(5):584–92. doi:10.1038/ncb1721

17. Diaz D, Recio JS, Weruaga E, Alonso JR. Mild cerebellar neurodegeneration of aged heterozygous PCD mice increases cell fusion of Purkinje and bone marrow-derived cells. Cell Transplant. 2012;21(7):1595–602. doi:10.3727/096368912X638900

18. Corish P, Tyler-Smith C. Attenuation of green fluorescent protein half-life in mammalian cells. Protein Eng. Dec 1999;12(12):1035–40. doi:10.1093/protein/12.12.1035

19. Ridder K, Keller S, Dams M, et al. Extracellular vesicle-mediated transfer of genetic information between the hematopoietic system and the brain in response to inflammation. PLoS Biol. Jun 2014;12(6):e1001874. doi:10.1371/journal.pbio.1001874

20. Kemp K, Gray E, Wilkins A, Scolding N. Purkinje cell fusion and binucleate heterokaryon formation in multiple sclerosis cerebellum. Brain. Oct 2012;135(Pt 10):2962–72. doi:10.1093/brain/aws226

21. Wiersema A, Dijk F, Dontje B, van der Want JJ, de Haan G. Cerebellar heterokaryon formation increases with age and after irradiation. Stem Cell Res. Nov 2007;1(2):150–4. doi:10.1016/j.scr.2008.02.001

22. Espejel S, Romero R, Alvarez-Buylla A. Radiation damage increases Purkinje neuron heterokaryons in neonatal cerebellum. Ann Neurol. Jul 2009;66(1):100–9. doi:10.1002/ana.21670

23. Agashiwala RM, Louis ED, Hof PR, Perl DP. A novel approach to non-biased systematic random sampling: a stereologic estimate of Purkinje cells in the human cerebellum. Brain Res. Oct 21 2008;1236:73–8. doi:10.1016/j.brainres.2008.07.119

24. Andersen BB, Gundersen HJ, Pakkenberg B. Aging of the human cerebellum: a stereological study. J Comp Neurol. Nov 17 2003;466(3):356–65. doi:10.1002/cne.10884

25. Cook AA, Fields E, Watt AJ. Losing the Beat: Contribution of Purkinje Cell Firing Dysfunction to Disease, and Its Reversal. Neuroscience. May 10 2021;462:247–261. doi:10.1016/j.neuroscience.2020.06.008

